# Repurposing of Anti-infectives for the Management of Onchocerciasis using Machine Learning and Protein Docking Studies

**DOI:** 10.1101/2023.02.26.530155

**Authors:** Cyril Tetteh, Bernice Ampomah, Mahmood B. Oppong, Michael Lartey, Paul Owusu Donkor, Kwabena F.M. Opuni, Lawrence A. Adutwum

**Author notes:** **Correspondence:** Lawrence A. Adutwum^1^.

## Abstract

The discovery of new drugs for the treatment of neglected tropical diseases (NTDs) is hampered by the lack of financial reward associated with their development. A combination of exploratory data analysis, machine learning (ML) and molecular docking studies were used to evaluate 58 anti-infective agents to identify those with the potential to be repurposed for the management of onchocerciasis, an NTD. Out of the 58 test drugs, 14 were predicted by at least five ML models to be useful in managing onchocerciasis. Molecular docking studies using glycine receptor subunit α-3, gamma-aminobutyric acid receptor subunit β-3 and glutamate-gated chloride channel of the 14 drugs showed binding affinities comparable to that of doramectin and pyrvinium which are known onchocerciasis drugs. Diminazine, trimetinib, triclabendazole, cridanimod, vandetinib and trametinib were the top agents showing high binding affinities from the molecular docking studies. The binding affinities of diminazine and cridanimod are similar to doramectin and pyrvinium which have demonstrated activity against onchocerciasis. The outcome of this study shows the potential of using these 14 drugs to manage onchocerciasis.

**Graphical Abstract:** 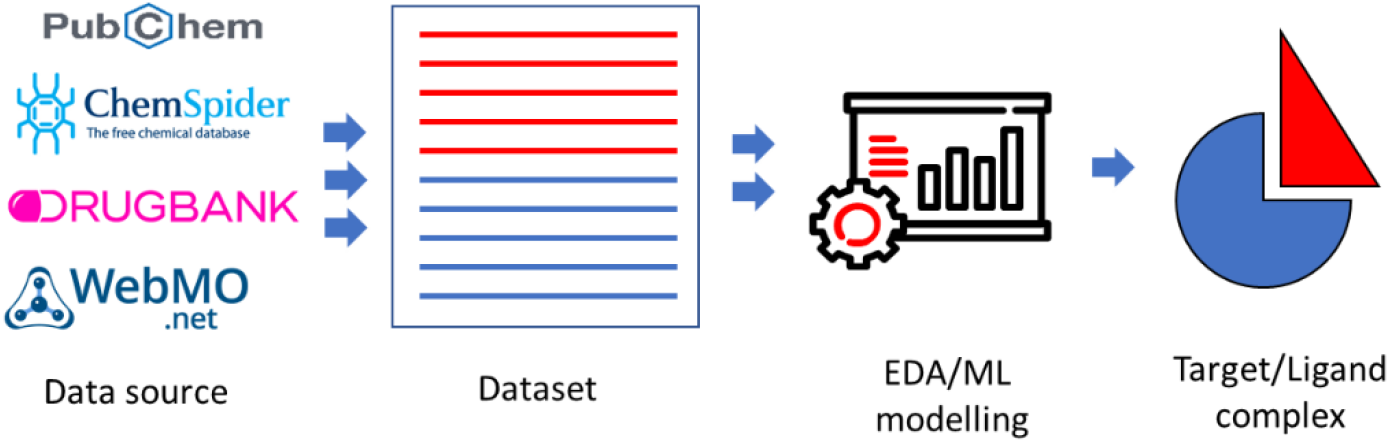

## 1. Introduction

Neglected tropical diseases (NTDs) are a group of diseases that affect low-income populations in the developing countries of Africa, Asia, and the Americas ^1–3^. Approximately one billion people, predominantly in developing countries living in remote rural areas, urban slums, or conflict zones, are at risk of NTDs ^3,4^. Onchocerciasis (river blindness or Robles’ disease) is a skin and eye infection caused by *Onchocerca volvulus*, a parasitic worm^5,6^. The disease is transmitted through the bites of infected blackflies, and symptoms of the disease include bumps under the skin, extreme itching, psoriasis, and loss of skin elasticity, among others. The most severe manifestations are ocular lesions that can progress to visual impairment and blindness^7^.

Ivermectin remains the main drug used to treat the disease^8–10^. Side effects such as neurotoxicity, central nervous system depression and liver toxicity have been reported^11,12^. Studies conducted in Ghana revealed that subjects repeatedly treated with ivermectin exhibited sub-optimal responses to the drug^13^. Even though several approaches are used to control the spread and the management of onchocerciasis, research on finding new therapeutic agents is not actively pursued^14^. Other drugs used in the past for managing onchocerciasis included diethylcarbamazine, suramin and moxidectin.^15–24^ Suramin use had to be discouraged due to the risk of optic atrophy^16^, while diethylcarbamazine is less effective even at high doses with frequent and severe side effects^25^. Pharmacokinetic studies on moxidectin show that it is an attractive long-acting therapeutic option for treating human onchocerciasis^31^ and was approved by the United States FDA in 2018^23,24^.

New drug discovery is confronted with many challenges including huge expenses, time, and the increasing stringency of regulatory agencies in evaluating the efficacy of new drug candidates^26–29^. Research into finding new chemotherapeutic agents for NTDs is highly unattractive because endemic areas are largely poverty-stricken. Due to the onerous nature, associated risk, and low return on research and development investments, there is limited interest in developing medications for treating NTDs including onchocerciasis ^28^.

Drug repurposing/repositioning is the process of finding new therapeutic indications for existing drugs reducing research cost on toxicities and activity^26,30–32^. This reduces the time and cost of drug discovery as information on doses and toxicities is already established. Repurposing of already existing drugs implies that research and costs associated with all pre-clinical work are also exempted^31,32^. As an alternative, it offers scientists and pharmaceutical companies an efficient strategy to identify novel targets and uses for already approved pharmaceutical agents. This provides the advantage of reducing the effort, time, and cost of every drug development stage for an area that is plagued by a need for affordable and effective therapies^32^. In fact, the latest drug moxidectin is a veterinary anthelminthic which has been repurposed for to manage onchocerciasis. Even though high throughput screening allows for the screening of several thousands of agents in a relatively short time, techniques that can weed out agents with low probability of being hits will be beneficial.

Recently, the use of ML methods to screen compounds in databases to find new indications have become popular^27,33–36^. Among others, concepts such as artificial intelligence, deep learning, re-enforcement learning have been used^33,34,37–40^. Halicin, an effective antibiotic was discovered using deep learning approaches^38^. ML methods have been used to identify drug candidates, which have been repurposed for managing diseases such as Alzheimer’s, cancers, and COVID-19^33,34,37,41–45^. Unfortunately, there is a dearth of literature on the use of ML methods to identify potential drug candidates that can be repurposed for NTDs such as onchocerciasis.

In this work, exploratory data analysis (EDA) and ML techniques were used to screen several anti-infective agents to identify potential candidates that may have activity against onchocerciasis. Subsequently, molecular docking studies were done on potential candidates to determine their binding interactions and binding affinities using targets that typically bind to drugs used for onchocerciasis.

## 2. Experimental Section

### 2.1. Dataset generation and Pre-processing

Three-dimensional (3D) structures of 602 anti-infective drugs were obtained from online repositories including DrugBank^46^, PubChem^47^, and ChemSpider^48^. Where the 3D conformers of the compounds were unavailable, their two-dimensional (2D) conformers were converted to their optimized 3D structures using WebMO (version 21.0)^49^. The web-based cheminformatics platform, online chemical modeling environment (OCHEM) and AlvaDesc^®^ (version 2.0.4) were used to generate molecular descriptors ^50–52^. This yielded a dataset matrix of 602 rows × 5,666 columns (rows × columns).

Descriptors with entries for less than 30% of the data were eliminated. This reduced the number of descriptors to 4,474. Of the 602 samples, 16 are/were used (or have demonstrated activity) in the management of onchocerciasis while 588 were not. The data thus comprised of two classes, which are onchocerciasis and non-onchocerciasis drugs which were labeled ***oncho***^**+**^ and ***oncho***^-^, respectively. The data was mean centered and auto scaled to unit variance.

### 2.2. Data Analysis

#### 2.2.1. Linear Discriminant Analysis (LDA) and Cluster Analysis (k-Means)

Using the scaled data, LDA was performed as a dimensionality reduction step using singular value decomposition solver. The significant uses of the drugs (12 indications) were used as the target. This was followed by cluster analysis using the ***k-means*** cluster algorithm to identify drugs like the ***oncho***^**+**^. The pairwise distance between the ***oncho***^**+**^ drugs and all other anti-infectives were determined. Five (5) anti-infectives closest to each onchocerciasis drug were selected and put into a test set. In total, 56 samples were captured in this group and were thus put into a test set. This reduced the training set data to 546 (i.e., 16 ***oncho***^**+**^, 530 ***oncho***^**-**^, 56 test drugs). Doramectin and pyrvinium, agents demonstrating activity against onchocerciasis were added to the test drugs to evaluate our proposed routine. Hence, the distribution of the data set for subsequent analysis became 14 ***oncho***^**+**^, 530 ***oncho***^**-**^, and 58 **test drugs** (including doramectin and pyrvinium).

#### 2.2.2. Class Imbalance and Machine Learning

The imbalance in the data, (14 ***oncho***^**+**^, 530 ***oncho***^***-***^) was corrected using the Synthetic Minority Oversampling Technique^53,54^ (SMOTE). A total of 516 artificial compounds were generated to balance the data. Thus, the final dataset for subsequent ML modeling was 1060 (530 ***oncho***^**+**^ & 530 ***oncho***^-^)

The dataset was split into 2/3 (707 drugs) and 1/3 (353 drugs) for training and validation sets, respectively. Ten supervised learning algorithms namely, Support Vector Machines (SVM), Random Forest (RF), Logistic Regression (LR), k-Nearest Neighbor (KNN) classification, Stochastic Gradient Boosting (SGB), Boosted Decision Trees (BDT), Naïve Bayes Classifiers (NBC), AdaBoost (ABC), Multi-Layer Perceptron (MLP), Gaussian Naïve Bayes (GNB) were used. These ML models were generated with the training set data using scikit-learn. Hyperparameter tuning was performed automatically to achieve the optimum model. The models’ performance was evaluated using their prediction accuracy as defined by Long *et.al*.^55^ on the external validation set. The models with prediction accuracies of at least 80% were used to evaluate the 58 test compounds.

#### 2.2.3. Protein Target Retrieval and Virtual Screening (Molecular Docking)

The binding affinity of test drugs predicted to have ***oncho***^**+**^ activities were evaluated using molecular docking studies. Four proteins identified as relevant to the management of onchocerciasis were used as targets. These were glycine receptor subunit α-3 (GRS-α-3, UniProt ID: O75311)^56^, gamma-aminobutyric acid receptor subunit β-3 (GABA-β-3, UniProt ID: P28472)^57^, Glutamate-gated chloride channel (GluCl, PDB ID: Q25634)^58^, and prostaglandin G/H synthase 1 (PGHS, UniProt ID: P23219)^59^. The PDB files for the crystal structures of these four proteins were downloaded from RCSB Database (https://www.rcsb.org/) and the SWISS-MODEL Repository (https://swissmodel.expasy.org/). The retrieved structures were cleaned up with Discovery Studio version 21.1.0 (BIOVIA, San Diego) ^60^ to correct issues associated with incomplete structures due to missing atoms or water, the presence of multimers and interaction partners of the receptor molecule.

Virtual screening was performed using the Python Prescription Virtual Screening tool (PyRx 0.8) ^61^ containing AutoDock Vina module ^62^. The prepared protein structure was fed into the PyRx tool along with the 3D conformers of the test drugs predicted to be ***oncho***^**+**^. The drug and protein molecules were converted to a PDBQT file using the AutoDock module of the PyRx tool. A grid box was then constructed to define docking spaces. The dimensions of the grid box were set along the *x, y*, and *z* axes at: 104.2, 102.5, 137.9Å for GABA-β-3, 130.9, 87.2, 126.6 Å for GRS-α-3, 154.1, 75.0, 122.7 Å for GluCl and 81.8, 69.9, 75.3 Å for PGHS. These parameters were set to encompass the entire 3D structure of each protein so that the ligand could freely move and rotate in the docking space. The 2D and 3D interactions between the protein-ligand were analyzed using Discovery Studio version 21.1.0 (BIOVIA, San Diego) ^60^.

## 3. Results and Discussion

The discovery of new drugs can be a long, tedious, and costly endeavour. Consequently, pharmaceutical companies invest in projects with higher chances of financial success. Thus, research into the discovery of some medications may be lacking even if they are needed by the population. The case is much worse for NTDs as the people in need of these drugs lack the financial capabilities to afford them. Drug repurposing presents a relatively cheaper route to discovering new drugs by finding new indications for already existing drugs. In this study, we explored EDA and ML techniques to identify anti-infectives that may have the potential to be repurposed for the management of onchocerciasis. To support the ML predictions, molecular docking studies were performed using known targets of drugs used to manage onchocerciasis to determine the interactions and binding affinities of the predicted agents.

ML algorithms benefit from the availability of large data sets. Unfortunately, fewer agents are used in the management of onchocerciasis leading to an imbalance in the dataset. With only 14 samples assigned to the ***oncho***^**+**^ class, the high number of descriptors relative to the sample size could lead to overfitting. Using a typical feature selection algorithm would not be ideal, hence, LDA was performed as a dimensionality reduction step.

Figure 1 (a and b) shows the LDA results using the major indications of the anti-infectives (Figure S1) as the output. The LDA latent variables (LDs) scatter plot for the first four LDs are shown in Figure S2 in the SI. The LDs in LDA are constructed to maximize the separation between the samples, i.e., significant uses. Subsequently, 11 LDs were determined to be ideal since they capture 99.77 % of the variability in the data.

**Figure 1.**
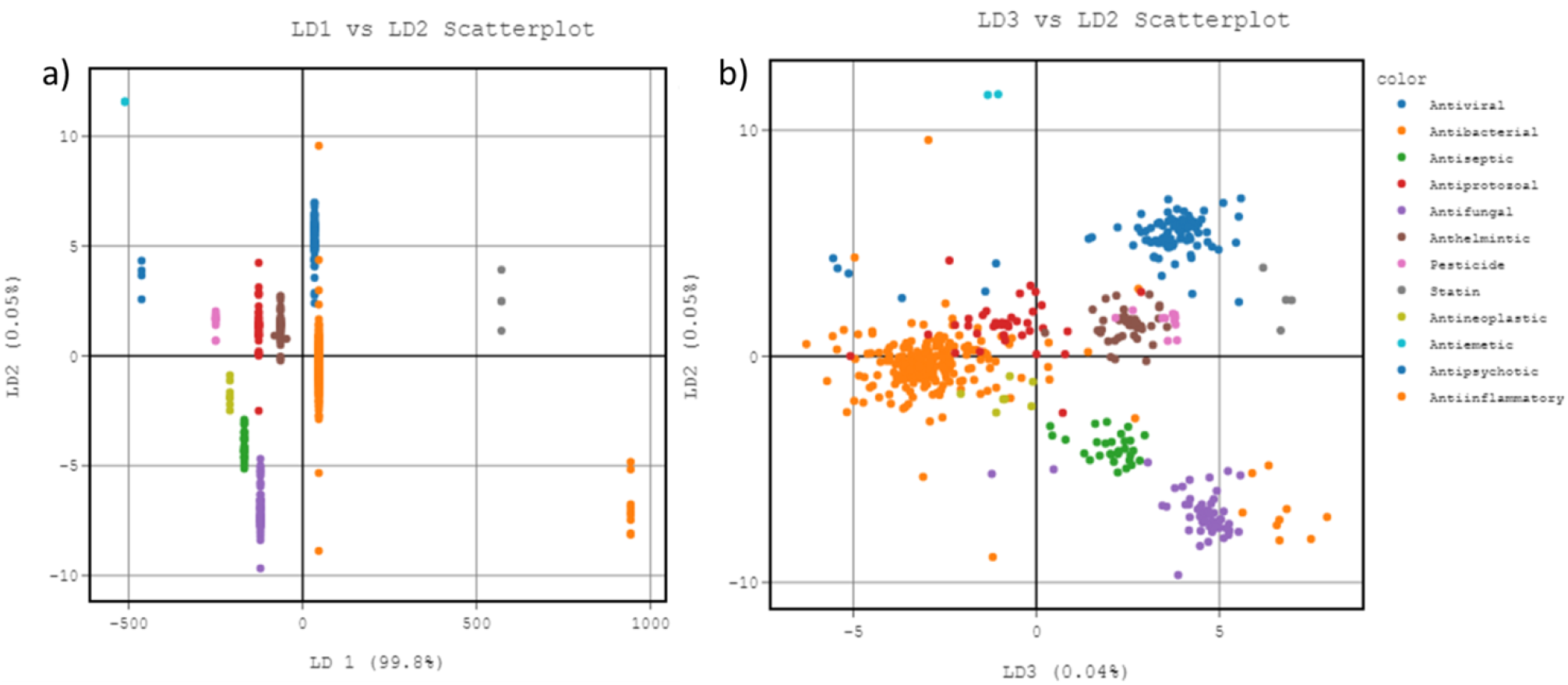
Visualizing the LDA model of anti-infective agents using the major indications/drugs (a) LD1 vs LD2 and b) LD3 vs LD2).

To find agents that could be repurposed for managing onchocerciasis from the data, an unsupervised learning approach, ***k-means*** cluster analysis was employed. In cluster analysis, drug samples that have similar properties/indications are expected to be grouped closer to each other. The ***k-means*** clustering algorithm was used due to its fast computation and general ability to produce tighter clusters^63,64^. Using the Folk Mallows and Rand Scores, an optimal ***k*** value of 11 was identified. The 11 clusters were just one less than the total number of indications of the drugs in the data set. Subsequently, the 14 ***oncho***^***+***^ samples were identified and the pairwise distance from these 14 drugs and the other samples were computed. The five (5) drugs that clustered closest to the ***oncho***^+^ drugs were selected. It is expected that their closer proximity to the ***oncho***^+^ drugs implied the likelihood to have similar therapeutic properties. A total of 70 drugs would be found in the test set, however, there were 56 drugs as some agents were found to be close to more than one ***oncho***^+^ drug. Figure S3 in SI is LD2 vs LD3 plot of the LDA model showing the location of ***oncho***^+^*(red)*, ***oncho***^-^ *(blue)* and test drugs *(orange)*. It must be highlighted here that doramectin and pyrvinium with known ***oncho***^***+***^ were added to investigate whether this approach would have identified them as potentially useful drugs^65,66^.

Generation of ML models with imbalanced data is not ideal as it leads to high model accuracies even if the models perform woefully in the minority class. The Imbalanced-learn SMOTE algorithm was implemented to generate samples to balance the data in the minority class. In total, 516 artificial ***oncho***^***+***^ samples were generated to balance the dataset. A plot of LD1 vs LD2 and LD2 vs LD3 for the dataset with and without the artificial samples can be found in Figure S4 in the SI.

ML models were generated using the balanced data as described earlier. Two-thirds of the data was used for the training set whiles a third was used for external validation. The following ML algorithms were used: SVM, RF, GNB, NB, KNN, LR, SGB, BDT, MLP and ADB. All these algorithms were used since one could not tell which would perform better for this dataset. Hyperparameter tuning was performed automatically. The model prediction accuracies on the external validation sets were used to evaluate the ML models. Figure 2 shows the accuracy of the ten different machine learning algorithms in predicting anti-infective agents as, specifically when tested on an external validation set of data. It can be seen here that with the same dataset and pre-processing, different algorithms give different prediction accuracies. Hence, valuable information may be lost if a single model is relied on. Consequently, the test drugs in this study were evaluated using an ensemble of the models. However, since all the models have different prediction accuracies, a threshold of at least 80% was selected. Seven (7) models met this criterion: SVM, RF, KNN, BDT, SGB, MLP and ADB. Subsequently, the seven (7) models were used to predict the onchocercidal activity of the 58 test drugs. The threshold prediction probability was set to a default value of 0.5. Thus, for a model, any drug with a *y* predicted ≥ 0.5 is deemed to have the potential to have onchocercidal activity. The number of drugs predicted by each algorithm is shown in Figure S5 in the SI. Figure S5 shows KNN predicted 38 of the test drugs as having the potential to be used for onchocerciasis with SGB predicting the least 13. Figure 3a-g shows the *y* predicted psrobabilities of the 58 test drugs. The results indicate varying predicted probabilities for the test drugs by different ML algorithms. These variations in predicted probabilities for different algorithms are expected due to the various approximation in the objective functions used in solving ML problems. Thus, using several algorithms allows us to truly identify which drugs have true potential. Due to the variation in the number of predicted samples by each ML model, drugs selected by at least 5 of the 7 ML models were selected for molecular docking studies.

**Figure 2.**
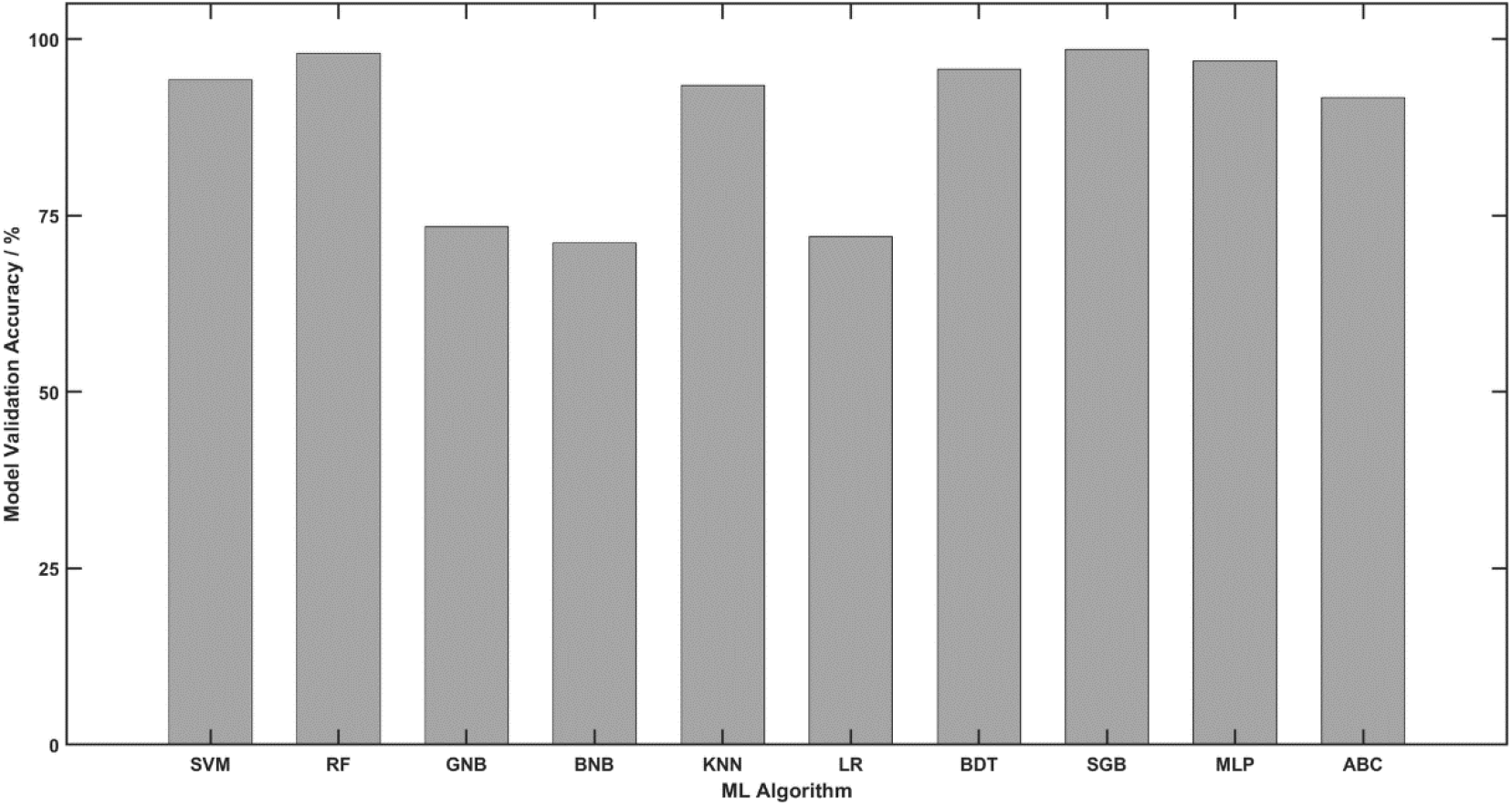
The model prediction accuracies of the ten (10) machine learning algorithms using the external validation set data.

**Figure 3.**
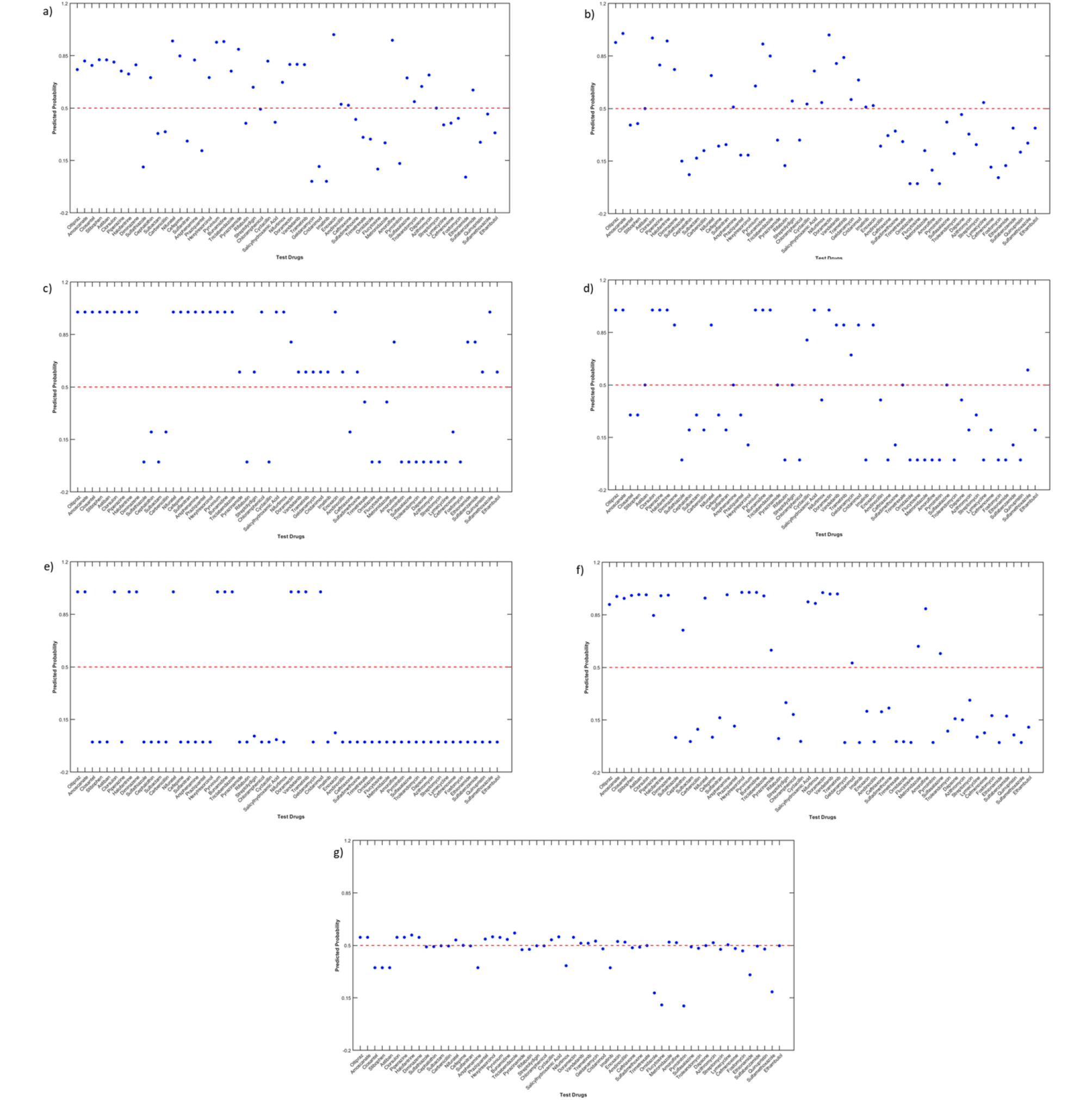
The y-predicted probability plots of the 16 test samples (including pyrvinium and doramectin) for the top seven best performing ML algorithms based on the model prediction accuracy on the external validation set (a) SVM, b) RF, c) KNN, d) BDT, e) SGB, f) MLP and g) ADB. The red dashed line represents the 0.5 predicted probability value above which a drug is classified as ***oncho***^***+***^.

**Table S1** in SI shows the list of the test drugs and their predictions by each of the 7 selected models. Sixteen (16) drugs representing 27.59 % of the test data were predicted to be ***oncho***^***+***^. It is noteworthy that onchocercidal drugs doramectin and pyrivinium were predicted as ***oncho***^***+***^ by all 7 models. This affirms that a combination of EDA and ML truly has the potential to aid in the repurposing of anti-infectives for NTDs.

### Molecular Docking Studies

The drug of choice in the management of onchocerciasis is ivermectin. Its antiparasitic activity is attributed to its binding to ivermectin-sensitive glutamate-gated chloride channel (GluCl) and glycine receptor subunit α-3 (GRS-α-3)^67–69^. In mouse models, evidence shows that the anticonvulsant effect of Ivermectin is due to its binding to gamma-aminobutyric acid receptor subunit β-3 (GABA-β-3)^69^. Diethylcarbamazine, another anti onchocerciasis agent is thought to be involved in sensitizing the micro-filariae to phagocytosis through its interaction with Prostaglandin G/H synthase 1 (PGHS) in the arachidonic acid metabolic pathway^70^. Thus, for any of the predicted drugs to have some activity against onchocerciasis it must bind to some, if not all, of these target proteins. Thus, the binding interaction with these targets can provide useful information to further boost the confidence in the drugs. The binding affinity, Kd measures the interaction between two molecules - a macromolecule (e.g., protein//target) and a ligand (drug) ^71^. Low Kd indicates higher affinity whiles higher Kd indicates lower affinity.

The binding interaction between these four receptors and the 16 drugs was performed as described earlier. The results of these experiments are shown in Figure 4. It is evident from Figure 4 that all the drugs predicted by the ML algorithms have binding interactions with all the protein targets studied. It is not surprising that our evaluation drugs, doramectin and pyrvinium are the 1^st^ and 2^nd^ drugs in all targets except GRS-α-3 where pyrvinium was 3^rd^. The detailed binding interactions of the top three for each target excluding the evaluating drugs were examined.

**Figure 4.**
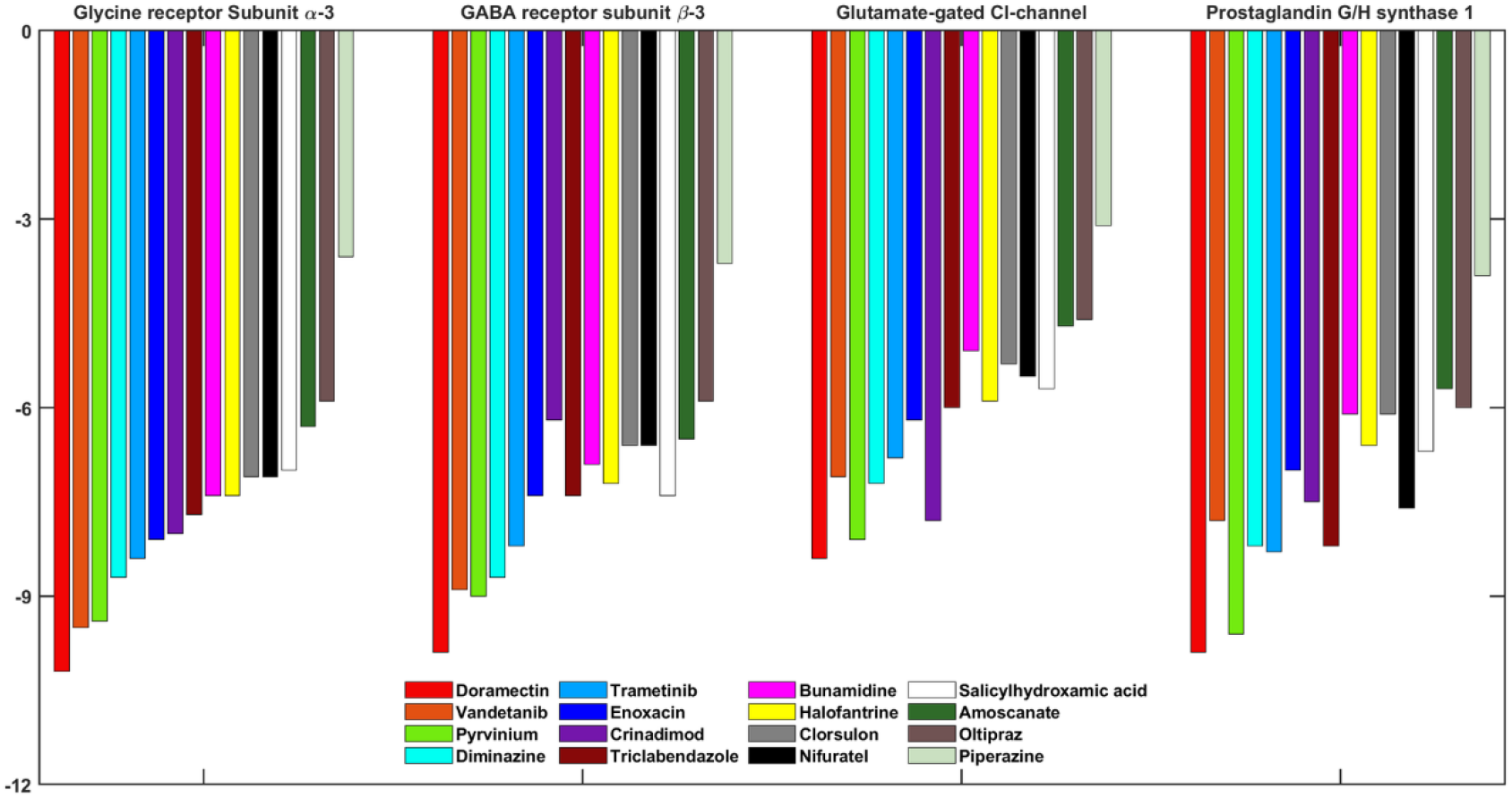
The binding affinity (Kd) of the 16 test samples (including pyrvinium and doramectin) for the GluCl, GRS-a-3, GABA-b-3 and PGHS receptors.

The binding interaction of the top three drugs for GRS-α-3 namely, vandetinib, trimetinib, and diminazine are show in Figure 5. Vandetanib showed a binding affinity of -9.5 kcal/mol to GRS-α-3 which is higher than pyrvinium, the evaluation drug. Vandetanib forms a hydrogen bond and an anion-pi interaction with the ALA26 and ASP86 residues while Trametinib forms pi-sigma bond and a halogen interaction with the ALA254 and PRO250 residues. Diminazine forms four hydrogen bonds with SER82, TYR161, THR113 and ARG27 as well as a pi-pi T-shaped interaction with PRO10 residues of GRS-α-3.

**Figure 5.**
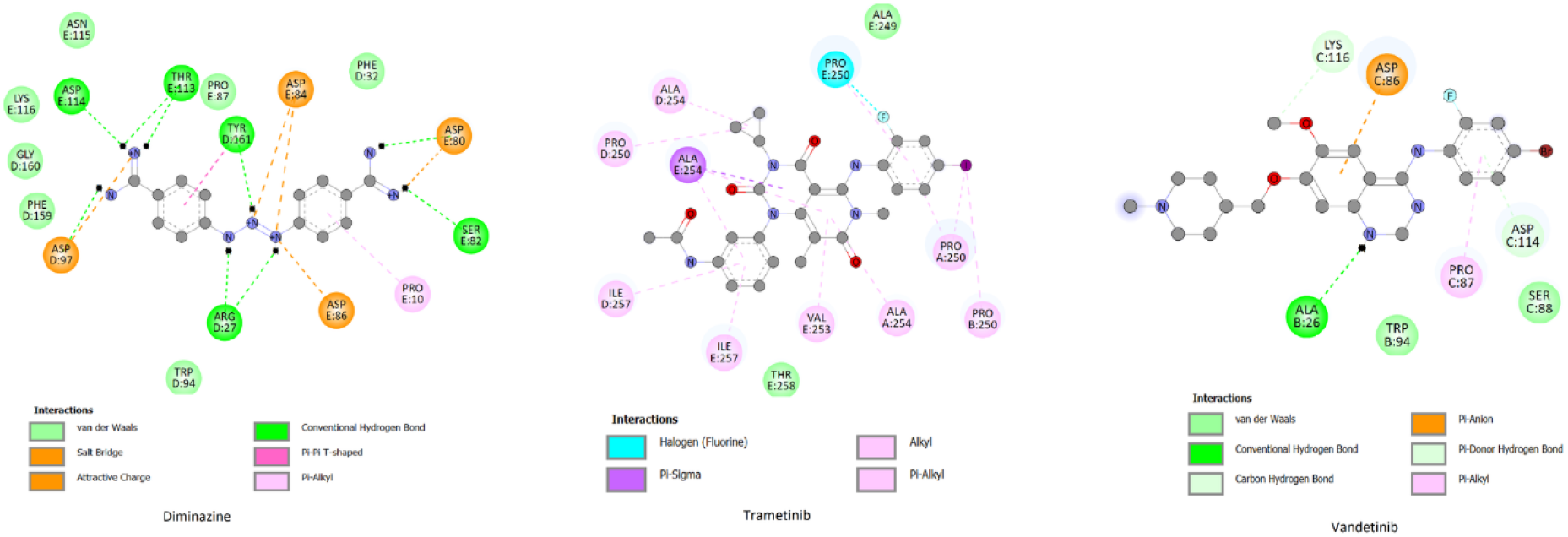
The binding interaction between GRS-a-3 and Diminazine, Trametinib and Vandetinib.

It is interesting to note that vandetanib, diminazine and trametinib also turned to be the top three drugs that bind to GABA-β-3. From Figure 6, it can be seen that vandetinib forms three hydrogen bonds with ASP48, VAL50 and VAL53, a pi-sulphur bond with MET137 and a halogen interaction with GLU182. Vandetanib also forms a pi-alkyl bond with LYS274, a residue found to link with pyrivinium through pi-donor interaction. Diminazine forms a pi-pi stacked interaction with TYR62, a pi-sigma interaction with ALA201 and several hydrogen bonds with ASN41, TYR97, SER156, TYR157, ALA174 and LYS173. Trametinib forms a pi-pi stacked interaction with PHE306, a pi-pi T-shaped interaction with TYR304 and two hydrogen bonds with ARG428 and TYR304.

**Figure 6.**
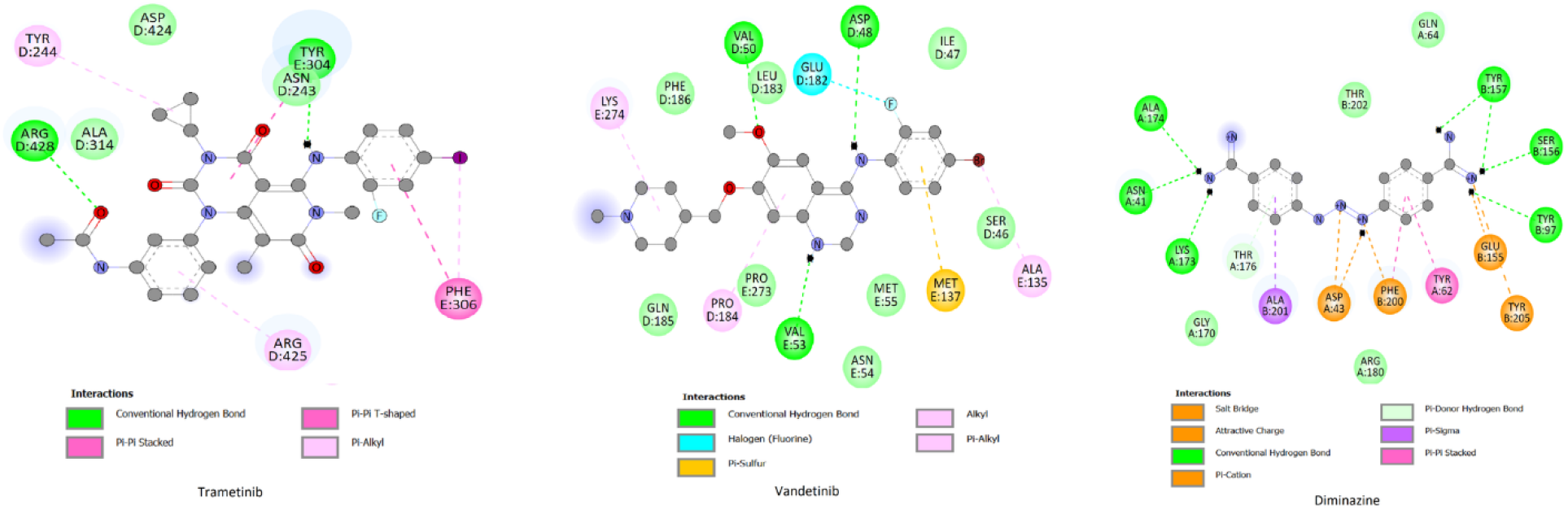
The binding interaction between GABA-β-3and Diminazine, Trametinib and Vandetinib

The binding affinities for cridanimod, diminazine and vandetanib with GluCl were -7.8, -7.2 and -7.1 kcal/mol respectively. Cridanimod is a small molecule functionally related to acridone and shown to have antineoplastic adjuvant activity. The compound interacts with GluCl via hydrogen bonding at GLU25, ASN45, ASN46, and GLN103; pi-anion at GLU25 and pi-alkyl at VAL40 and ARG41. Diminazine forms two hydrogen bonds at PRO216 and ILE70, pi-pi stacked at TYR254 similar to pyrivinium. However, there was shown to be an unfavorable donor-donor interaction with SER217. Vandetanib forms a halogen bond with GLN258 and pi-cation interaction with LYS305 (Figure 7).

**Figure 7.**
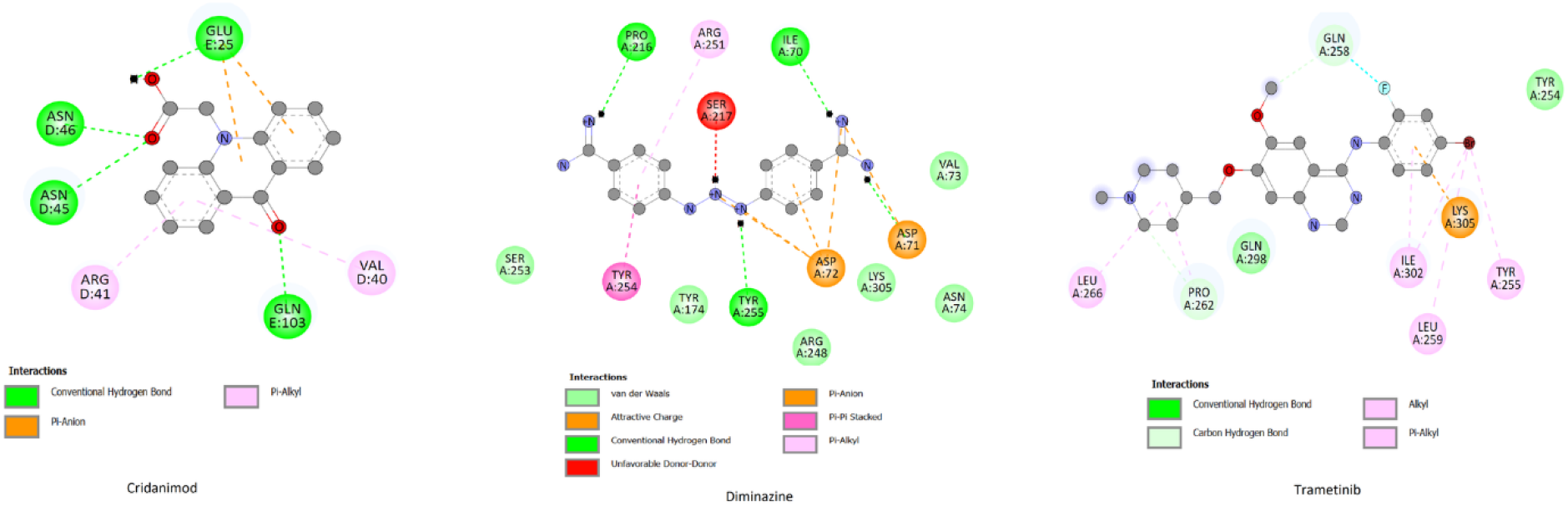
The binding interaction between GluCl and Diminazine, Trametinib and Cridanimod

In the case of PGHS, diminazine, trimetinib and triclabendazole demonstrated the highest binding. Detailed binding interactions are shown in Figure 8. Triclabendazole is a narrow-spectrum anthelminthic with activity against *Fasciola* and *Paragonimus spp*. It shows a binding affinity of -8.2 kcal/mol and interacted with PGHS via hydrogen bonding with the amino acid residues CYS41 and CYS47 as well as halogen bonding and pi-alkyl interaction with CYS47 and PRO153 respectively. Trametinib forms one hydrogen bond with ARG120, a pi-sigma bond with VAL119 and a pi-cation interaction with ARG83 while Diminazine forms numerous hydrogen bonds with the residues TRP387, PHE210, MET391, ALA199, THR206, TYR385 and THR212 which was also found to interact with Doramectin through hydrogen bonding. Diminazine also forms three pi-interactions with PHE210, HIS388 and ALA202 and a pi-cation interaction with HIS386.

**Figure 8.**
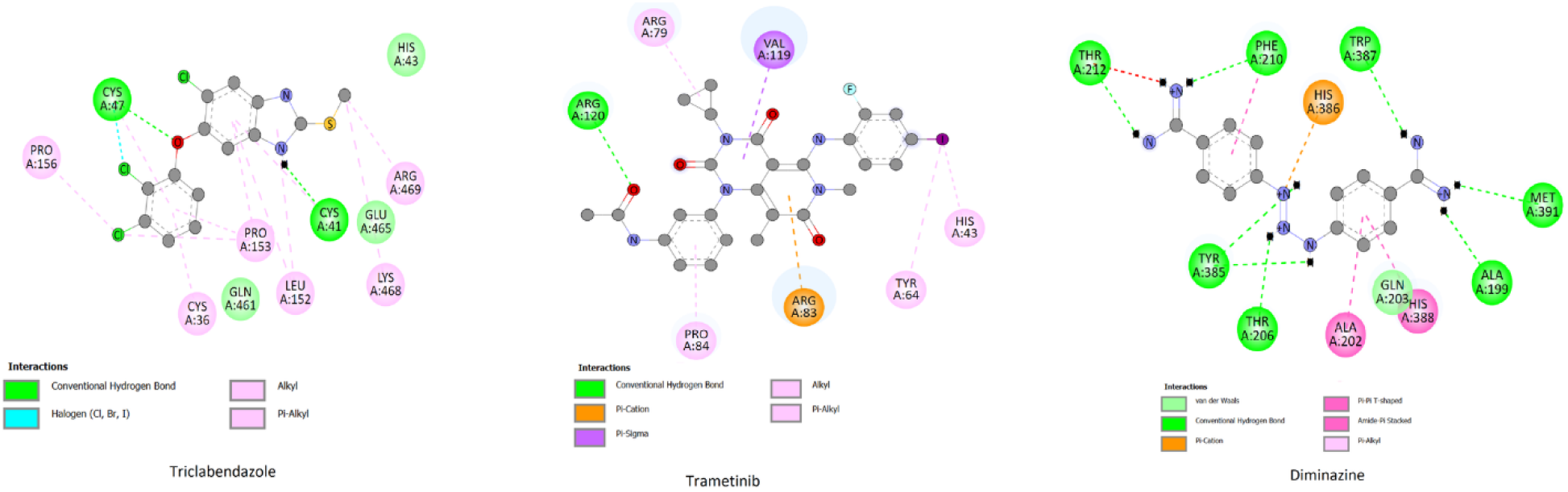
The binding interaction between PGHS and Triclabendazole, Trametinib and Diminazine.

## 4. Conclusion

Using exploratory data analysis, machine learning and molecular docking studies, 14 anti-infectives have been identified to possess the potential to be repurposed for the management of onchocerciasis. Out of these 14 drugs, six showed strong binding interactions comparable to drugs used in the management of onchocerciasis. These six are diminazine, trimetinib, triclabendazole, cridanimod, vandetinib and trametinib. Of these drugs, diminazine exhibits the highest potential to be used for onchocerciasis. These compounds could also serve as lead compounds in the discovery of new and improved therapies for onchocerciasis.

The techniques and routines implemented here can be applied to other research areas to find new indications for existing drugs.

## 5. Conflict of Interest

*The authors declare that the research was conducted in the absence of any commercial or financial relationships that could be construed as a potential conflict of interest*.

## 1 Data Availability Statement

The datasets for this study can be found in the https://github.com/lawrenceadutwum/ntdrepurpose.

## Notes

### Competing Interest Statement

The authors have declared no competing interest.

